# Deep learning models for RNA secondary structure prediction (probably) do not generalise across families

**DOI:** 10.1101/2022.03.21.485135

**Authors:** Marcell Szikszai, Michael Wise, Amitava Datta, Max Ward, David H. Mathews

**Affiliations:** Department of Computer Science & Software Engineering, The University of Western Australia, Perth, WA, Australia; The Marshall Centre for Infectious Diseases Research and Training, The University of Western Australia, Perth, WA, Australia; Department of Molecular and Cellular Biology, Harvard University, Cambridge, MA, USA; Department of Biochemistry & Biophysics, Center for RNA Biology, and Department of Biostatistics & Computational Biology, University of Rochester, Rochester, NY, USA

## Abstract

**Motivation:** The secondary structure of RNA is of importance to its function. Over the last few years, several papers attempted to use machine learning to improve *de novo* RNA secondary structure prediction. Many of these papers report impressive results for intra-family predictions, but seldom address the much more difficult (and practical) inter-family problem.

**Results:** We demonstrate it is nearly trivial with convolutional neural networks to generate pseudo-free energy changes, modeled after structure mapping data, that improve the accuracy of structure prediction for intra-family cases. We propose a more rigorous method for inter-family cross-validation that can be used to assess the performance of learning-based models. Using this method, we further demonstrate that intra-family performance is insufficient proof of generalisation despite the widespread assumption in the literature, and provide strong evidence that many existing learning-based models have not generalised inter-family.

**Availability:** Source code and data is available at https://github.com/marcellszi/dl-rna.

## 1 Introduction

Ribonucleic acid (RNA) molecules are extremely versatile polymers fulfilling numerous roles essential for life, including gene regulation and catalytic functions [1, 2]. Part of this versatility can be attributed to the structural diversity of RNA [3]. While chemically related to DNA, RNA often functions as a single strand. As a consequence, the molecules often fold back on themselves forming complex structures. It is well established that these folded configurations are of importance to the function of non-coding RNAs (ncRNAs) [4].

When discussing RNA, its structure is generally divided into a hierarchy of three levels. First, the foundation is the *primary structure*, which refers to the one-dimensional sequence of the molecule. The sequences are made up of a succession of nucleobases, represented by four letters: adenine (`A`), cytosine (`C`), guanine (`G`), and uracil (`U`). Next, the *secondary structure*, which refers to the set of canonical base pairings where bases are paired with one or zero other bases. For secondary structure, these pairs are formed by Watson-Crick base pairings (`A-U`, `G-C`) and by wobble `G-U` pairs. Finally—the last level generally considered— is the *tertiary* structure which refers to the three-dimensional structure and the additional interactions that mediate the structure. However, since the secondary heavily informs the tertiary structure [5, 6, 7], secondary structure is usually sufficient for developing some understanding of function.

Sequencing RNA molecules today is quick, inexpensive, and accurate [8]; however, determining their structure is not. While high-resolution experimental techniques—such as nuclear magnetic resonance spectroscopy, X-ray crystallography, and cryo-electron microscopy—exist, these methods tend to be expensive and time consuming. The contrast in the difficulty of determining sequence versus structure has created a sequence-structure gap, where there are vast amounts of sequenced RNA molecules without any known corresponding structure. In order to bridge this gap, significant effort has gone into developing algorithms to predict RNA structures computationally [4, 9, 7].

Broadly speaking, we can divide the secondary structure prediction methods into three categories: *homology modelling, comparative analysis*, and *de novo* methods. Methods that start with nothing but the sequence, often termed *de novo*, have the advantage of being effective for single sequences without a need for homologous sequences. However, these *de novo* methods are not always accurate and have well understood limitations [10]. These methods are generally implemented with dynamic programming based tools and often make use of an underlying thermodynamic model to determine the minimum free energy (MFE) structures [11, 12, 13, 14, 15]. In contrast, homology modelling and comparative analysis are more accurate, but require a set of homologous RNAs (and for homology modelling, their secondary structure). The sets of homologous RNA sequences are termed an RNA family [16]. These methods work by predicting a consensus structure that is conserved by evolution [17, 18, 19, 20].

In the last few years, a number of methodologies were published based on deep learning that report impressive results for RNA secondary structure prediction. However, many of these papers assess performance using *k*-fold cross-validation or simple train-test splits. We refer to these splits as *intra-family* (i.e. within-family), since there is no expectation that the families contained within the training and testing sets do not intersect. In contrast, we refer to splits where there is no such intersection as *inter-family* (i.e. between-family). Since the structure of RNA is highly conserved intra-family, performance derived from these metrics does not demonstrate generalisation to novel RNAs [21]. Homology modelling, and comparative analysis to an extent, is already well suited to the intra-family problem, and can not only determine the structure with high accuracy when used by domain experts, it can also provide other important insights about the molecule, such as its function [17]. Because of this, the practical use cases of machine learning models with poor inter-family performance are limited.

## 2 Materials and methods

### 2.1 Demonstrative model

#### 2.1.1 Basic concept

A common way to improve the performance of *de novo* tools is to utilise data from structure probing experiments. One example of such an approach is a technique by Deigan *et al*. [22], that supplements dynamic programming based methods via *selective 2’-hydroxyl acylation analysed by primer extension* (SHAPE) [23, 24], by which the experiment identifies nucleotides that are in more flexible regions of the secondary structure. SHAPE is an inexpensive probing experiment that scores the reactivity of each nucleotide in the RNA sequence. The reactivities found through SHAPE can be used to construct pseudo-free energy change terms for each nucleotide via the function,

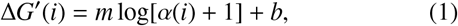

where *α*(*i*) is the SHAPE value for base *i*; *m* and *b* are free parameters, and log is the natural logarithm. Then Δ*G*′ is added as a free energy change term to each base pair stack involving nucleotide *i* in the *de novo* dynamic programming algorithm to improve predictive performance.

Our methodology looks to computationally mimic data from structure probing experiments, and ultimately construct pseudo-free energies that can be utilised by existing algorithms. We use this approach to construct a simple demonstrative model which shares many similarities with current learning-based efforts. We then show that our demonstrative model performs significantly better than existing dynamic programming based techniques for intra-family predictions. Finally, we show that our model performs poorly for inter-family predictions, demonstrating that intra-family performance does not necessarily generalise to inter-family cases.

Beyond our demonstrative model, we benchmarked or otherwise analysed several existing machine learning models for secondary structure prediction. See Section 2.4 for more details.

#### 2.1.2 Network architecture

We implemented a convolutional neural network (CNN) for extracting per-nucleobase pairing probabilities from RNA sequences with the aim of constructing pseudo-free energies to improve secondary structure prediction performance. The extracted pairing probabilities are then converted to pseudo-free energies and fed to RNAstructure [25] version 6.3, which makes use of a conventional dynamic programming algorithm to find the minimum free energy (MFE) structures.

The architecture of the neural network is made up of two blocks, a convolution block and a fully connected block. The convolution block is comprised of two one-dimensional convolution layers, with 256 filters each. The length of the kernels is 3, with strides of 1, and no dilation is applied. Each convolution layer is followed by a rectified linear unit (ReLU) activation layer. Spatial dropout [26] is applied between the convolution layers with a dropout rate of 0.25. The fully connected block is comprised of two fully-connected layers, with 512 neurons each. Dropout is applied between the layers with a dropout rate of 0.25. The first activation is once again ReLU, and the final layer is followed by sigmoid activation,

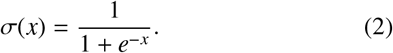

The network is trained with the Adam optimiser (*γ* = 0.001, *β*_1_ = 0.9, *β*_2_ = 0.999, *ϵ* = 10^−8^) [27] using binary cross-entropy loss and in mini-batches of 256. Early stopping is applied with a patience of 5.

#### 2.1.3 Encoding

For our demonstrative model, the input nucleotide sequences are one-hot encoded as two-dimensional matrices, where the nucleobases are represented by column vectors of size 4 and a sequence is the concatenation of these vectors. For example, a simple sequence `UCG…AC` is encoded as,

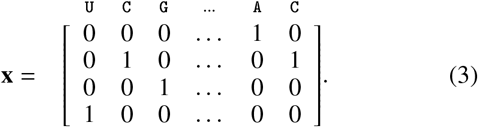

The target structures’ *shadows* are encoded as row vectors, classifying whether a particular base is paired or unpaired (without reference to the base pairing partner). RNA secondary structures can be represented by dot-bracket notation, where unpaired nucleotides are represented by ‘.’ characters, and paired nucleotides are represented by matches parentheses. Opening brackets indicate the 5’-nucleotide in a pair, and the matching closing brackets indicate the 3’-nucleotide in the pair.

These dot-bracket formatted structures can be easily converted to a structure’s shadow for our demonstrative model. For example, the simple sequence-structure pair from Figure 1 is encoded as,

**Figure 1:**
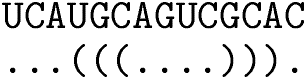
Example of a simple sequence-structure pair in dot-bracket format.

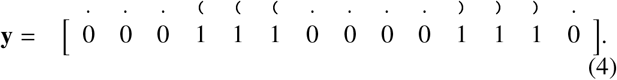

Both the sequences and structures are zero-padded at the 3’ ends, to have shape (4 × 512) and (1 × 512), respectively.

#### 2.1.4 Pseudo-free energy calculation

Since our neural network is designed to identify paired bases, *ŷ*(*i*) ≈ 1 indicates a nucleotide is predicted as likely to be base Preprint – Deep learning models for RNA secondary structure prediction (probably) do not generalise across families paired, while *ŷ*(*i*) ≈ 0 indicates a nucleotide is predicted as likely to be unpaired, where *ŷ*(*i*) is the predicted shadow of the structure at base *i*. This differs from SHAPE, where values close to zero increase pairing likelihood, and values far from zero decrease it. Because of this, we applied the transformation,

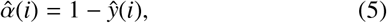

to our predicted values to produce SHAPE-like scores. These scores can then be used via the pseudo-free energy equation by Deigan *et al*. [22] (Equation 1) to improve the performance of the dynamic programming based MFE folding algorithms. Note that this transformation is the equivalent of redefining the pseudo-free energy equation as,

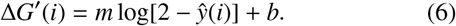

We performed grid-search on the slope *m*, and intercept *b* of Equation 6 to refit the model for the values extracted by our neural network. The search was performed on the validation set, and an 11 × 11 grid of slope and intercept values was investigated via a reduced version of the parameter space used by Deigan *et al*. [22]. Because the training and validation sets contain RNA sequences from the same families, overfitting these pseudo-free energy parameters may also be a concern. Creating non-intersecting training and validation sets was found to be problematic due to the limited number of families available in the dataset. Therefore, to address the inter-family generalisability of the computationally found pseudo-free energies, we explored the behaviour of the thermodynamic nearest neighbour model with small, uniform pseudo-free energy changes applied. These small free energy nudges are completely general, so they eliminate any underlying bias, and allow us to investigate whether uniform changes to the model affect the performance of MFE folding differently across families. For each sequence in our data set, we performed folds using RNAstructure [25] with pseudo-free energy change terms Δ*G*′ using parameters: *α*(*i*) = 0, *m* = 0, and *b* = {− 1.00, − 0.98, …, 0.98, 1.00}, totalling to 101 folds per sequence. That is, we apply the pseudo-free energy change term from Equation 1 with *m* = 0, so our pseudo-free energy nudges are given by,

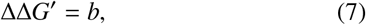

across the entire sequence. RNAstructure’s default tenths precision was increased to hundredths for improved resolution.

### 2.2 Data sets

Our experiments call for a large number of reliably known sequence-structure pairs, diverse in families. The data set used for this purpose is ArchiveII [28]. This dataset contains 3974 RNAs, across tRNAs [29], Signal Recognition Particle (SRP) RNAs [30], telomerase RNAs [31], 5S rRNAs [32], 16S rRNAs, 23S rRNAs [33, 34], tmRNAs [35], Group I [33, 34] and II Introns [36], and RNase P RNAs [37].

Most folding algorithms have polynomial time complexities *O*(*n*^*k*^) with *k* ≥ 3 [38], and the algorithm employed by RNAstructure is *O*(*n*^3^) [39]. Similarly, many of the learning-based models also suffer from significantly slower training and predictions for longer sequences. Because of this, we filter out 109 sequences longer than 512 nt to limit the runtime of our experiments, reducing our data set to 3865 RNAs. See Table 1 for a count of RNAs in each family after filtering.

**Table 1:** Breakdown of RNA families in ArchiveII after filtering.

| Family | Mean Length | # |
| --- | --- | --- |
| 5S rRNA | 119 | 1283 |
| SRP RNA | 180 | 918 |
| tRNA | 77 | 557 |
| tmRNA | 366 | 462 |
| RNase P RNA | 332 | 454 |
| Group I Intron | 375 | 74 |
| 16S rRNA<sup>1</sup> | 317 | 67 |
| Telomerase RNA | 438 | 35 |
| 23S rRNA<sup>1</sup> | 326 | 15 |
| Mean | 281 |  |
| Total |  | 3865 |

### 2.3 Train-test split

To assess intra-family performance, we perform *k*-fold cross-validation with *k* = 5 on our entire dataset. We note that despite its wide use in literature, it is our opinion that this type of train-test split cannot be used to assess the generalisation of machine learning models for RNA secondary structure prediction. We stress that this metric is used only to demonstrate the ease of achieving high accuracy for the intra-family case.

For benchmarking inter-family performance, we perform *family-fold cross-validation*, such that one family is left out for testing per cross-validation fold. The motivation behind this is to measure the models’ performance on novel RNAs that do not belong to a known family. This eliminates most of the homology to the training set, providing a fair measure of performance against other *de novo* tools.

In both cases, for early stopping and grid-search, we use a validation set which is a 10% randomly selected subset of the training set.

### 2.4 Existing models

We benchmarked or otherwise analysed several machine learning models for secondary structure prediction with a focus on investigating inter-family vs. intra-family performance. The models considered were: DMfold [41], RPRes [42], CROSS [43], E2Efold [44], SPOT-RNA [45], MXfold2 [46], and UFold [47] (Table 2).

**Table 2:** Recent papers that used machine learning for RNA secondary structure prediction. Inter-family and intra-family columns indicate the splitting methodology used in the paper, while the re-trained column indicates whether we have successfully re-trained the model on our dataset. Attempts were made to re-train every model, however, many do not publish training methodology or could not be re-trained for another reason. See Section 3.2, Section 4.2, and the Supplementary Information for a detailed discussion on this.

| Name | Authors | Year | Intra-family | Inter-family | Re-trained |
| --- | --- | --- | --- | --- | --- |
| CROSS | Delli Ponti *et al.* | 2017 | ✓ | ✗ | ✗ |
| DMfold | Wang *et al.* | 2019 | ✓ | ✗ | ✗ |
| SPOT-RNA | Singh *et al.* | 2019 | ✓ | ✗ | ✗ |
| E2Efold | Chen *et al.* | 2019 | ✓ | ✗ | ✗ |
| RPRes | Wang *et al.* | 2021 | ✓ | ✗ | ✗ |
| MXfold2 | Sato *et al.* | 2021 | ✓ | ✓ | ✓ |
| UFold | Fu *et al.* | 2021 | ✓ | ✓ | ✓ |

Where possible, we re-trained networks using family-fold cross-validation and benchmarked them for more generalised performance. In cases where we were unable to re-train the network, we provide evidence that the training/testing split does not appropriately consider RNA homology. All mentioned papers address intra-family performance, usually with simple *k*-fold cross validation, and in many cases wrongly conflate it with inter-family performance. The inter-family case is seldom mentioned, except by Sato *et al*. [46] and Fu *et al*. [47].

### 2.5 Benchmarking

To assess performance, we followed prior practice of calculating sensitivity and PPV [48]. We calculated the F_1_ score as the harmonic mean of positive predictive value (PPV) and sensitivity (SEN). Pairs were allowed to be displaced by one nucleotide position on either side so that for base pair `(i)-(j)` both `(i`±`1)-(j)` and `(i)-(j`±`1)` are considered valid [48]. Additionally, we performed two-tailed paired t-tests for statistical testing [48], considering *p* ≤ 0.05 significant.

Further, to assess the raw performance of our demonstrative model, we calculated the area under the curve of the receiver operating characteristic (AUC) for each RNA family. The receiver operating characteristic curve is constructed by plotting the sensitivity for predicting base pairing vs. the false positive rate (1 - specificity) at different threshold values. This metric allows us to measure how well our model can capture the secondary structure’s *shadow*. Specifically the AUC can be interpreted as the probability that our model can correctly distinguish between a paired and an unpaired nucleotide, so we can gain a better understanding of how well our classifier performs prior to any pseudo-free energy calculations.

## 3 Results

### 3.1 Demonstrative model

#### 3.1.1 Grid-search

Our experiments show that the behaviour of the thermodynamic nearest neighbour model is different across families for constant pseudo-free energy nudges (Equation 7) between − 1.0 and 1.0 kcal/mol (Figure 2). Effectively, a uniform nudge with negative value increases the stabilities of canonical pairs and nudges of positive value decrease the stabilities of canonical pairs. Much of these differences could be explained by noise, however the dramatic deviation as ΔΔ*G*′ → − 1 suggests that even under completely generalised inputs, such as these ΔΔ*G*′ nudges, the predictive performance of families is not uniformly affected. As an example, for some families like telomerase RNAs, the degradation in the negative region is much more extreme than in others like RNase P RNA. The differences are less clear when looking at ΔΔ*G*′ → +1, but are still present, especially when contrasting certain pairs like 23S RNA and tmRNA.

**Figure 2:**
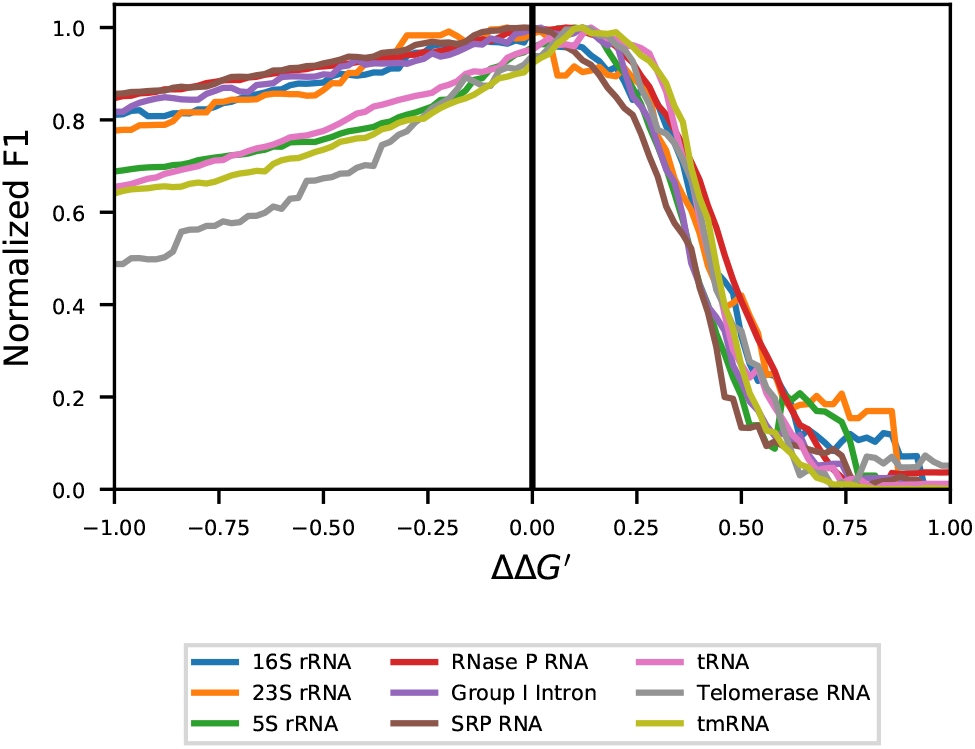
Comparison of the effect of ΔΔ*G*′ nudges between families. The mean of all sequences in each family is calculated across the ΔΔ*G*′ values. F_1_ scores have been normalised (min-max scaled) to account for the differences in underlying secondary structure prediction performance between families.

As expected, no ΔΔ*G*′, significantly improves performance across all families simultaneously. However, at least one region where the nudges improve performance can be found for all families with the exception of 16S rRNA. The region ΔΔ*G*′, ∈ [0.04, 0.10] improves the F_1_ score of 5S rRNA, RNase P RNA, tRNA, telomerase RNA, and tmRNA – although not necessarily significantly. Regions with significant improvements are [0.06, 0.14] for 5S rRNA, [0.20, 0.26] for tRNA, and [0.02, 0.22] for tmRNA. The performance of the remaining families: Group I, 16S RNA, 23S RNA, and SRP RNA is degraded within [0.04, 0.10] with significantly worse performance for 16S RNA and SRP RNA.

The grid-search for the slope (*m*) and intercept (*b*) free parameters of Equation 6 revealed that while the optimal values differ from those found with real SHAPE experiments [22, 49], the region of well-performing parameters is also present for generated SHAPE-like values (Figure 3). Our small pseudo-free energy nudges showed that even inherently general changes to the existing thermodynamic nearest neighbour model do not affect the families uniformly, so overfitting the slope and intercept values to our training set is likely. To minimise the possible impact of overfitting these parameters towards families present in the training set, we elected to use *m* = 1.8 kcal/mol and *b* = − 0.6 kcal/mol, as found by Hajdin *et al*. [49] for experimentally-generated SHAPE data. We expect that these parameters themselves are general and any overfitting to families is a result of the underlying generated SHAPE-like values.

**Figure 3:**
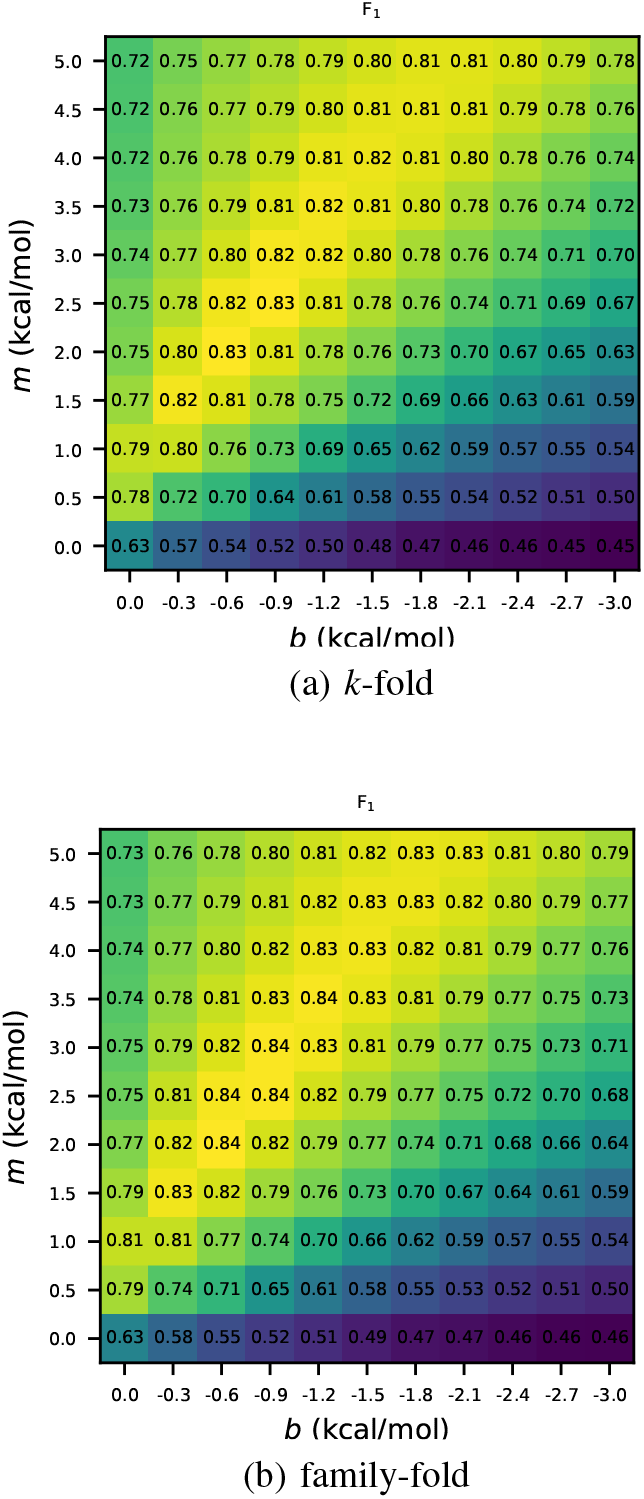
Results of the slope *m* and intercept *b* grid-search for the pseudo-free energy Equation 1 free parameters. A separate grid-search was done for *k*-fold and family-fold cross validation. Note the wider band of the optimal region for the simpler *k*-fold model, likely seen due to the strength of inter-family performance.

#### 3.1.2 Intra-family vs. inter-family performance

Our demonstrative deep learning model (Section 2.1) shows improvements in F_1_ score across most families (except Group I Intron, and 23S rRNA), when benchmarked using *k*-fold cross-validation, over RNAstructure’s baseline scores (Table 3). This confirms that our simple model is able to trivially improve intra-family predictions over traditional dynamic programming MFE algorithms. Note that this is true even with our conservatively chosen pseudo-free energy free parameters, and the relatively high AUC values across all families suggest that more aggressive optimisation would likely yield even better results. Most of this can be attributed to the similarity of structures within particular families. While we see significant improvements (Supplementary Information) using *k*-fold cross-validation, this is not true for family-fold cross-validation. The F_1_ score is degraded across all families when compared to the baseline RNAstructure predictions, and AUC values are also significantly worse (see Supplementary Information) for all families when compared to *k*-fold cross-validation. It seems reasonable to expect that this is simply explained by training for too many epochs since there is strong family overlap between the training and validation sets (so early stopping does not prevent overfitting). However, examining the per-epoch performance of any split (Figure 4) suggests that this is not the case, and the model never shows signs of generalising for inter-family cases.

**Table 3:** Performance of the demonstrative model separated by RNA family. F_1_ score refers to the performance of secondary structure prediction and AUC refers to the performance of predicting the structures’ shadow via deep learning. The baseline is RNAstructure for free energy minimization without the deep learning input. Both *k*-fold and family-fold models are included.

| Family | # | Baseline | $k$-fold | | family-fold | |
| --- | --- | --- | --- | --- | --- | --- |
| | | $F_1$ | AUC | $F_1$ | AUC | $F_1$ |
| 5S rRNA | 1283 | 0.63 | 0.95 | 0.94 | 0.72 | 0.46 |
| SRP RNA | 918 | 0.64 | 0.88 | 0.81 | 0.73 | 0.50 |
| tRNA | 557 | 0.80 | 0.97 | 0.97 | 0.79 | 0.65 |
| tmRNA | 462 | 0.43 | 0.82 | 0.64 | 0.68 | 0.41 |
| RNase P RNA | 454 | 0.55 | 0.81 | 0.66 | 0.71 | 0.48 |
| Group I Intron | 74 | 0.53 | 0.73 | 0.53 | 0.72 | 0.49 |
| 16S rRNA | 67 | 0.58 | 0.77 | 0.60 | 0.72 | 0.48 |
| Telomerase RNA | 35 | 0.50 | 0.76 | 0.61 | 0.68 | 0.45 |
| 23S rRNA | 15 | 0.73 | 0.79 | 0.68 | 0.73 | 0.54 |
| Total | 3865 |  |  |  |  |  |
| Mean |  | 0.60 | 0.83 | 0.72 | 0.72 | 0.50 |

**Figure 4:**
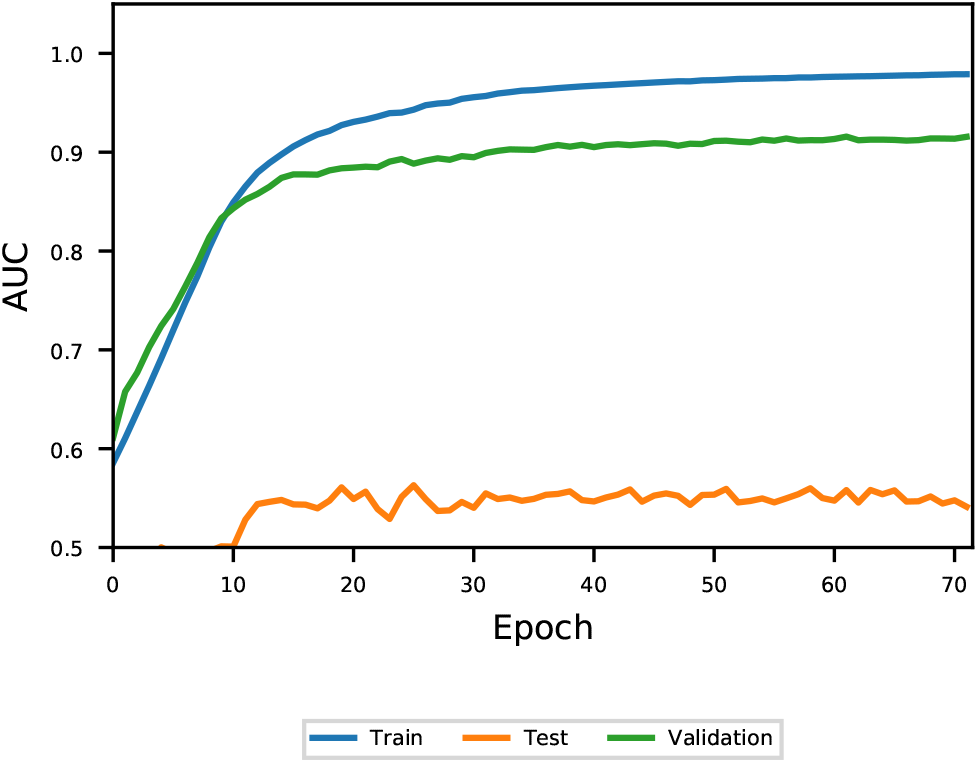
Performance of family-fold testing on our demonstrative model. The training set is comprised of all families except 5S rRNA, the validation is a 10% split of the training set, while the testing set is 5S rRNAs. Note the consistently poor performance of the testing set throughout.

**Figure 5:**
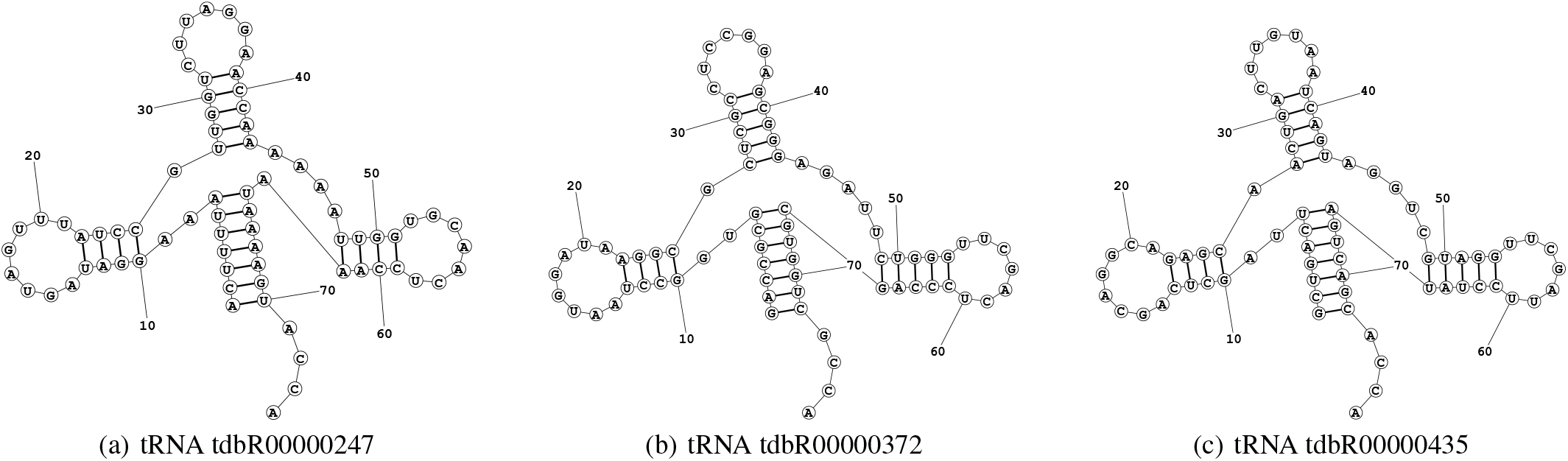
Secondary structure of three tRNAs. Despite relatively low sequence identity (<60%), their secondary structures appear nearly identical. Many machine learning model benchmarks fail to separate these RNAs between the training and testing sets, causing significant overlap.

The 36% difference (F_1_ = 0.72 to F_1_ = 0.50, Table 3) in performance between intra-family vs. inter-family cases is strong evidence that *k*-fold cross-validation is insufficient for bench-marking deep learning methods for RNA secondary structure prediction.

### 3.2 Existing models

In order to evaluate how well existing models generalise, we attempted to re-train all of their networks using family-fold cross-validation to benchmark inter-family performance (Table 4). Unfortunately, many of these tools do not publish their source code, particularly for training. Further, we were unable to retrain a number of models with public source code due to bugs in the code, which in some cases prevented us from being able to run their tools at all. Please see Table 2, Section 4.2 and the Supplementary Information for a detailed discussion on this.

**Table 4:**
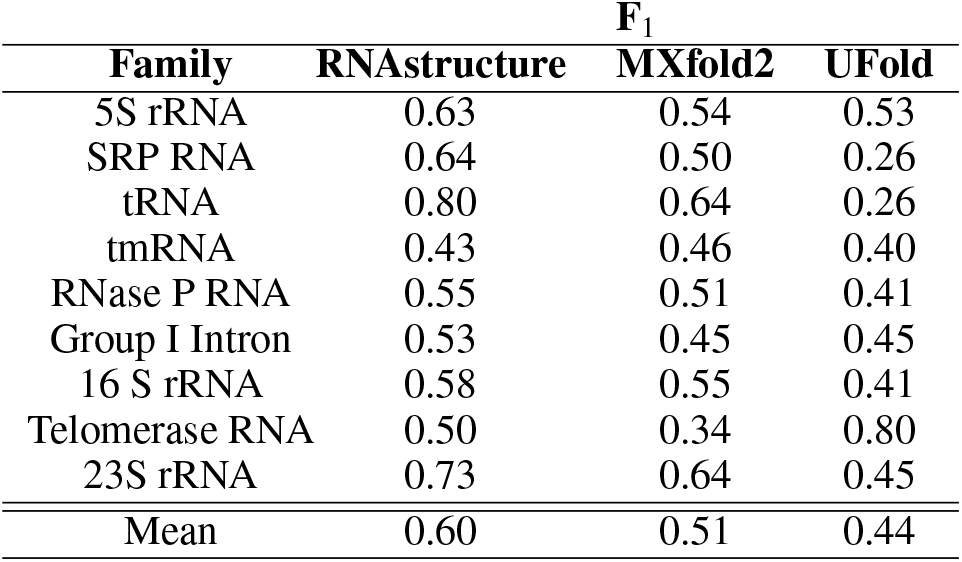
Performance of family-fold cross-validation on MX-fold2 and UFold.

## 4 Discussion

### 4.1 Demonstrative model

First, our results indicate that pseudo-free energy change terms affect RNA families differently. We propose that it is possible to overfit the estimation of these parameters to specific structures or families. For example, after adapting Deigan *et al*. [22]’s equation (Equation 1) for alternate SHAPE-like values, refitting the parameters *m* and *b* requires careful consideration. This is especially true for any learning-based models attempting to improve RNA secondary structure prediction, since they require significant data to train already and may suffer from underlying overfitting issues. Our pseudo-free energy nudges have no inherent bias towards any family, so it is possible that any model that is not completely general may suffer even more dramatically.

We were able to make use of RNAstructure’s default parameters, found by jackknife resampling [49] across several families, which has successfully eliminated issues with generalisability for real SHAPE experiments. These parameters were within the optimal region for our extracted SHAPE-like probing information, so the performance degradation is minimal. However, it is worth noting that intra-family performance can be further improved by optimising *m* and *b* for the new distributions of *α* for base paired and unpaired nucleotides. Unfortunately, this can exacerbate issues with overfitting to the intra-family case even further.

In the case of the learned base pairing probabilities, or more generally, in the case of all hyperparameter optimisation tasks, creating unbiased training and validation sets is a challenge. After early-stopping for our demonstrative model, we were able to look at the performance of the test set per epoch, and observe that we were not over-training. However, in a model that is able to generalise, our validation set would be insufficient for any sort of hyperparameter optimisation. While the use of a training, validation, and testing set is commonplace in machine learning tasks, for RNA homology, the overlap of the families within these sets is the most important consideration. Even considering sequence identity or similarity measures is not enough, as structure is so highly conserved amongst families. Ideally, in order to do hyperparameter optimisation for learning-based models fairly, there can be no intersection between the families in the training, validation, and testing sets. This can be difficult when the number of accurately known RNA structures in the dataset is fairly small, and covers only a few families.

Second, our demonstrative model was able to achieve high AUC in the intra-family case, however, completely failed to generalise when it came to the inter-family case. This alone is evidence that it is insufficient to show good performance on intra-family predictions since it is, at the very least, possible to construct a model that does not work for practical applications and achieves high intra-family performance. Metrics like *k*-fold cross-validation as used for the demonstrative model, do not address RNA homology to any extent, since they do not address the intersection between families.

We suggest that benchmarking of learning-based methods for RNA secondary structure prediction be done by family-fold cross-validation in order to minimise the possibility of overfitting and accurately measure generalisation. Previous work by Rivas *et al*. [21], focusing on generative models instead of deep learning models, also supports our conclusions. For the purpose of fair benchmarking, a split is provided by this paper that attempts to minimise homology between training and testing sets as much as possible; however, it should be noted that the relatively small number of sequences in ArchiveII [28] means that we expect generalisation to this dataset to be difficult. It should be noted that the split on families reduces the concerns on homology, but does not completely eliminate all concerns about generalization. tmRNA, for example, is tRNA-like and mRNA-like [50]. Therefore, the tRNA-like features could overtrain a model which cross validation with tRNA would not reveal.

### 4.2 Existing models

For any machine learning model, an unbiased split of training and testing data is essential for benchmarking performance. In the case of biological data, this means considering the homology between these sets carefully in order to eliminate their overlap. Many current studies in RNA secondary structure prediction, especially those using learning-based models, do not appropriately address RNA homology. While it may be sufficient in many bioinformatics applications to consider sequence identity or sequence similarity, in the case of RNA, structure is strongly conserved amongst families – often much more than sequence. Because of this, it is possible (and highly probable) to create splits where despite considering sequence similarity, near identical structures are present in both the training and testing data sets. Below is a breakdown of the training/testing split methodologies used by existing methods.

#### 4.2.1 CROSS & RPRes

Both Computational Recognition of Secondary Structure (CROSS) [43], and RPRes [42] are methods that attempt to recreate SHAPE experiments *in silico*, sharing many similarities with our demonstrative model. Unfortunately, no source code is provided for CROSS, and as such, we were unable to re-train their model on our dataset. While the authors of RPRes do publish the source code on Github, we were unable to re-train their network. See the Supplementary Information for more details.

Neither paper sufficiently addresses concerns regarding poor inter-family generalisation. Both models are evaluated by training on one dataset at a time (PARS yeast, PARS human, HIV SHAPE, icSHAPE, and high-quality nuclear magnetic resonance spectroscopy / X-ray crystallography structures) and testing on all others one-by-one. With this methodology, there is no guarantee, or indeed expectation, that the secondary structures in the datasets do not overlap. According to Delli Ponti *et al*. [43] “[n]egligible overlap exists between training and testing sets” [43] with Jaccard indices *<* 0.002 between each pairs of datasets, where *Jaccard*(*S* _1_, ∩ *S* _2_) = (*S* _1_ ∩ *S* _2_) */* (*S* _1_ ∪ *S* _2_) for sequences *S* _1_ and *S* _2_. This addresses sequence similarity, but does not comprehensively address inter-family cases.

#### 4.2.2 DMfold

While the authors of DMfold [41] do publish the entire source code on Github, we were unable to re-train their network. See the Supplementary Information for more details.

The train/test split methodology used by DMfold produces sets which heavily overlap families. After using their packaged tools for generating the splits, we found that all testing families were covered in the training set, without any consideration to RNA homology whatsoever. In this case, the training set contained 2111 RNAs, with 957 5S rRNAs, 437 tRNAs, 377 RNase P RNAs, and 340 tmRNAs, while the testing set contained 234 RNAs, with 102 5S rRNA, 49 tRNAs, 45 RNase P RNAs, and 38 tmRNAs. This set contains many identical, or nearly-identical structures between the training and testing sets, with a mean minimum tree edit distance of 14.16, compared to the 134.99 of our family-fold cross-validation splits. See the Supplementary Information for more details.

#### 4.2.3 E2Efold

While the authors of E2Efold [44] do publish the entire source code on Github, we did not re-train their network due to high memory requirements. However, other recent publications have already pointed out E2Efold’s poor inter-family performance, reporting F_1_ scores as low as F_1_ = 0.036 [46, 47] on the bpRNA-new dataset.

The original E2Efold benchmarks use stratified sampling, generating train/test splits which heavily overlap families. The training set, based on RNAStralign [20], contained 24,895 RNAs, with 9325 16s rRNAs, 7687 5S rRNAs, 5412 tRNAs, 1243 Group I Introns, 379 SRP RNAs, 431 tmRNAs, 360 RNase P RNAs, and 28 telomerase RNAs. The first testing set, based on RNAStralign once again, contained 2825 RNAs, with 1150 16s rRNAs, 879 5S rRNAs, 504 tRNAs, 136 Group I Introns, 53 SRP RNAs, 61 tmRNAs, 37 RNase P RNAs, and 5 telomerase RNAs. The second training set, based on ArchiveII [28], explicitly only contained families that overlap with the RNAStralign dataset.

#### 4.2.4 MXfold2 & UFold

Our re-training of MXfold2 [46] and UFold [47] with family-fold cross-validation indicates that the models do not generalise well to inter-family performance. However, as previously pointed out, it could be argued that our tests are particularly hard due to the small number of families in our dataset.

Sato *et al*. [46] did address inter-family performance using their own bpRNA-new dataset for which they reported positive results. To address inter-family performance, the model is trained on bpRNA-1m, a dataset derived from Rfam 12.2 [51]. The model is then tested on bpRNA-new, which is derived from a newer version of Rfam (14.2) [31]. Newly discovered and novel RNA families are extracted from Rfam 14.2 making up the bpRNA-new testing set.

Since this testing set does not share any families with the training set, we expect that good performance on this split provides reasonable evidence for generalisation. It should be noted however, that these results are still less robust than our proposed family-fold cross-validation, since secondary structures amongst families are often similar in Rfam, particularly within “clans”, which are “group[s] of families that either share a common ancestor but are too divergent to be reasonably aligned or a group of families that could be aligned, but have distinct functions” ^2^. For example, the SRP clan is divided into 9 separate families. There are clear homologs within this clan, such as, for example, `Fungi_SRP` and `Metazoa_SRP`. Because of this, to firmly address inter-family generalisation, Rfam families should also be split by clan. Additionally, reporting of the performance should always be broken down by family to provide context about generalisation.

UFold [47] applies the same testing methodology as MXfold2, and reports similarly positive results, although according to self-reported metrics [47] on the bpRNA-new dataset, both tools are outperformed by Eternafold [52], a multitask-learning-based method that uses a crowdsourced RNA design data set [53] to train a model, and Contrafold [54], a statistical learning method that uses conditional log-linear models.

#### 4.2.5 SPOT-RNA

SPOT-RNA [45] does not provide the source code for training, or the ability to re-train the model. As such, we were unable to evaluate it on our dataset.

Singh *et al*. [45] initially pre-trained the model on bpRNA-1m [51], and then applied transfer learning to train, validate, and test on a small set of 217 high-resolution structures. Both the pre-training and training sets are separated from the testing set by filtering based on 80% sequence-identity, and BLAST-N [55] is used to address homology with an e-value cut-off of 10. While better than relying solely on a sequence-identity cutoff, BLAST-N itself is a sequence similarity based metric, and does not address the secondary structure of the RNAs, meaning that this split cannot be considered inter-family. Indeed, the self reported improvement of 20% (F_1_ = 0.49 to F_1_ = 0.58) over the dynamic programming based RNAfold [13], becomes a 3% (F_1_ = 0.62 to F_1_ = 0.60) [46] deterioration when benchmarked on MXfold2’s bpRNA-new dataset [46] of novel families.

## 5 Conclusion

Our results show that a basic convolutional neural network can be used to construct pseudo-free energies to improve secondary structure prediction for intra-family cases. We proposed the more rigorous testing methodology of family-fold cross-validation, which along with our model was used to demonstrate that intra-family performance does not guarantee generalisation to inter-family cases. We argued that *k*-fold cross validation is an unsuitable method for benchmarking deep learning RNA secondary structure prediction models. Finally, we used these findings as evidence that many recent publications wrongly conflate intra-family with inter-family results, and that this results in inflated self-reported accuracy.

## Footnotes

1 16S rRNA and 23S rRNA are split into independent folding domains. [40]

2 https://docs.rfam.org/en/latest/glossary.html#clan

