## Supplementary Information for "Deep learning models for RNA secondary structure prediction (probably) do not generalise across families"

Preprint, compiled March 21, 2022

**Marcell Szikszai** [1,*], **Michael Wise** [1,2], **Amitava Datta** [1], **Max Ward** [1,3], **and David H. Mathews** [4]

[1]Department of Computer Science & Software Engineering, The University of Western Australia, Perth, WA, Australia
[2]The Marshall Centre for Infectious Diseases Research and Training, The University of Western Australia, Perth, WA, Australia
[3]Department of Molecular and Cellular Biology, Harvard University, Cambridge, MA, USA
[4]Department of Biochemistry & Biophysics, Center for RNA Biology, and Department of Biostatistics & Computational Biology, University of Rochester, Rochester, NY, USA

**5 Demonstrative model results** **7**

**6 $\Delta\Delta G'$ nudges** **8**

### 1 Demonstrative model grid-search

#### *1.1 k-fold*

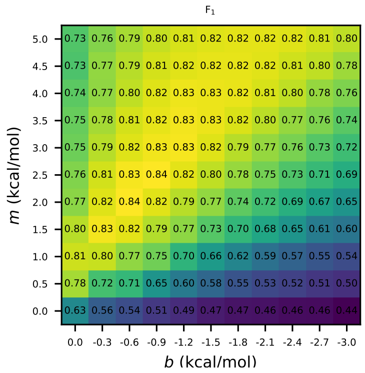

(a) $k = 0$

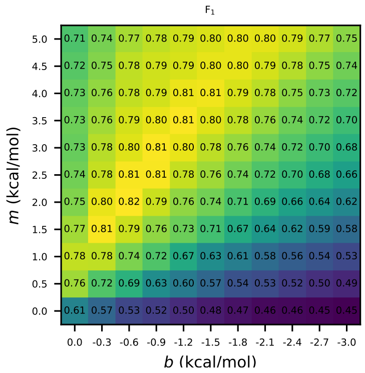

(b) $k = 1$

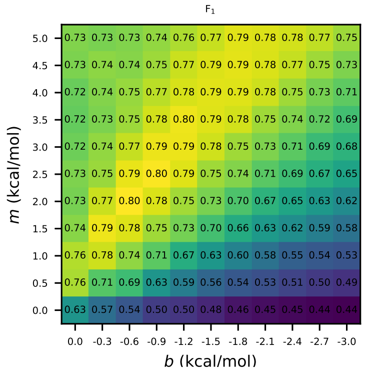

(c) $k = 2$

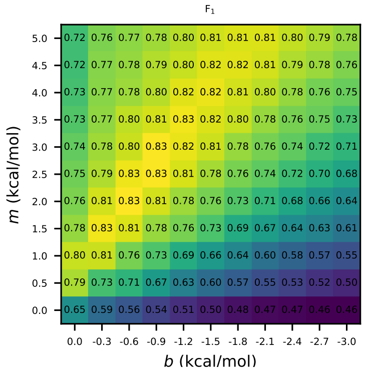

(d) $k = 3$

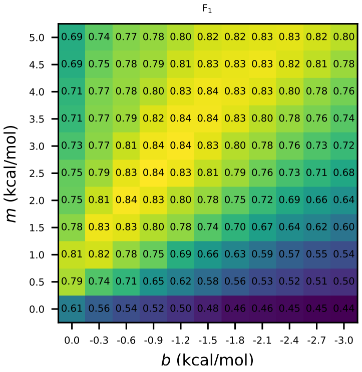

(e) $k = 4$

Figure 1: Results of the slope $m$ and intercept $b$ grid-search for the pseudo-free energy Equation 1 free parameters for $k$-fold cross-validation, broken down by $k$.

#### 1.2 Family-fold

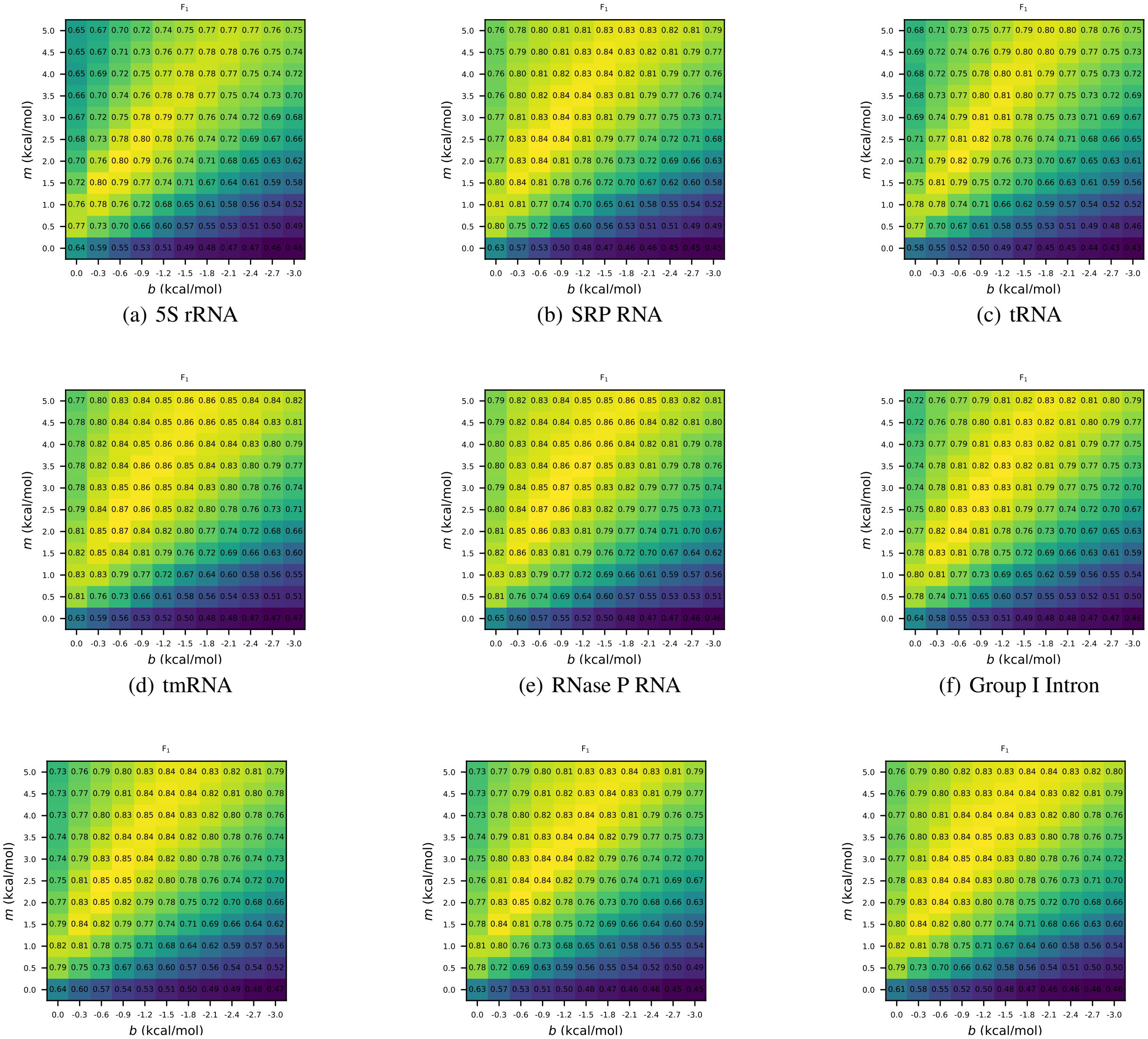

(a) 5S rRNA (b) SRP RNA (c) tRNA

(d) tmRNA (e) RNase P RNA (f) Group I Intron

(g) 16S rRNA (h) Telomerase RNA (i) 23S rRNA

Figure 2: Results of the slope $m$ and intercept $b$ grid-search for the pseudo-free energy Equation 1 free parameters for family-fold cross-validation, broken down by family.

### 2 Demonstrative model AUC vs. Epoch

#### 2.1 *k-fold*

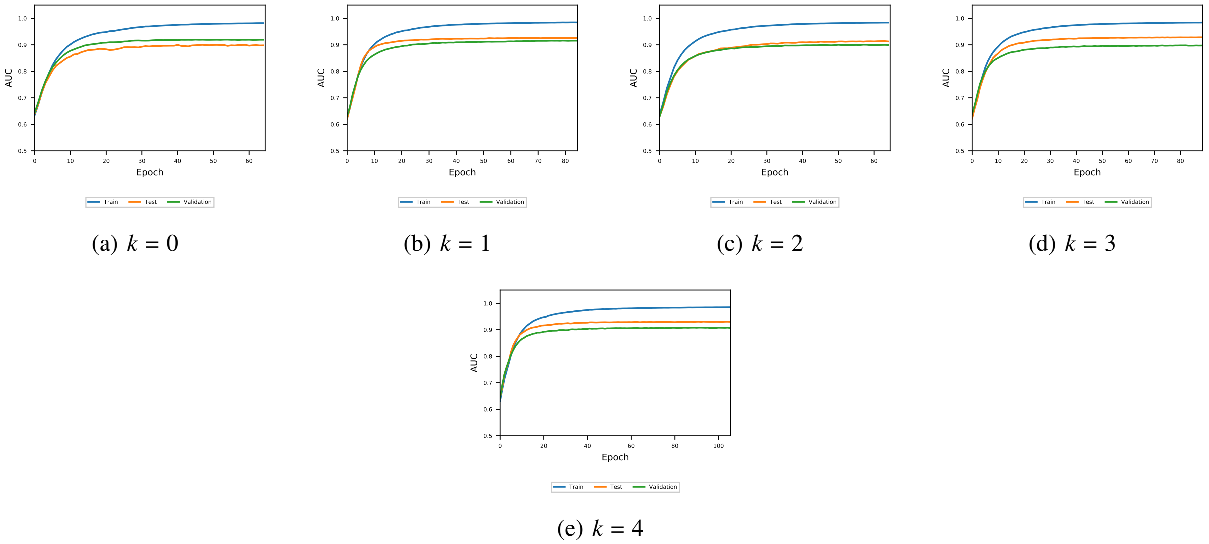

(a) $k = 0$ (b) $k = 1$ (c) $k = 2$ (d) $k = 3$ (e) $k = 4$

Figure 3: Performance of $k$-fold training, validation, and testing sets on our demonstrative model vs. training epoch.

#### 2.2 *Family-fold*

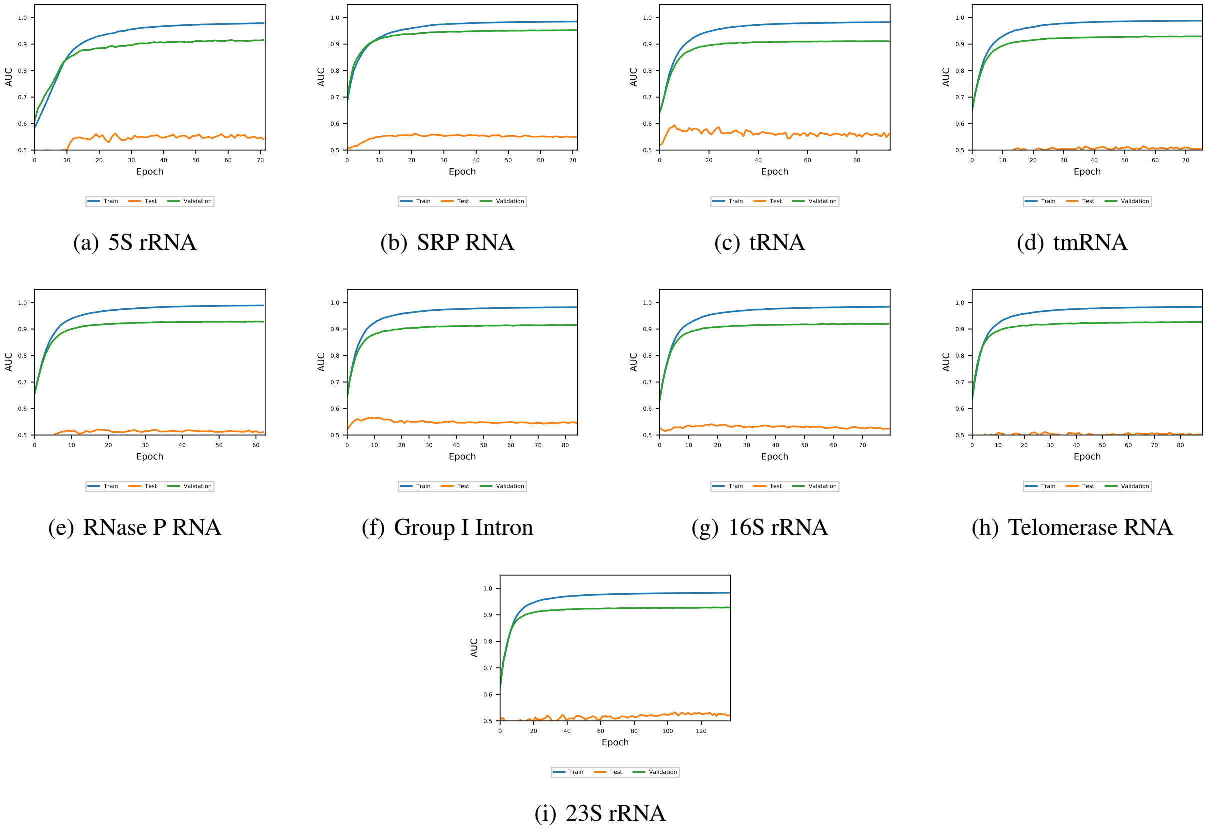

(a) 5S rRNA (b) SRP RNA (c) tRNA (d) tmRNA (e) RNase P RNA (f) Group I Intron (g) 16S rRNA (h) Telomerase RNA (i) 23S rRNA

Figure 4: Performance of family-fold training, validation, and testing sets on our demonstrative model vs. training epoch.

### 3 Training reproducibility

##### *CROSS*

No source code or training methods are provided for CROSS, and as such, we were unable to re-train their model on our dataset.

##### *3.1 DMfold*

The authors of DMfold publish the entire source code on Github, however, we were unable to re-train their network. The published Git repository contains bugs that make it impossible to reproduce their methods, or meaningfully debug their code. To attempt to run the program, we used Python 3.5.2 and Tensorflow 1.5.0 as specified in the provided Software Guide.

We have opened an issue on Github[1] detailing the problems we ran into trying to re-train their model, but received no response from the authors.

##### *3.2 SPOT-RNA*

No source code or training methods are provided for SPOT-RNA, and as such, we were unable to re-train their model on our dataset.

##### *3.3 E2Efold*

Source code is published, and training is reproducible, although the memory requirements are high (176GB+).

We contacted the authors via email (sent 2021-08-10), and they were helpful in assisting with re-training their method. However, as the MXfold2 [1] and UFold [2] papers have already benchmarked E2Efold for inter-family performance, we did not feel it was necessary to benchmark on our dataset.

##### *3.4 RPRes*

The authors of RPRes do publish the entire source code on Github, however, we were unable to re-train their network as per their instructions. Attempting to run RPRes out of the box throws a `RuntimeError: DataLoader worker exited unexpectedly` exception. There are very limited instructions provided on how to run the code, and no `requirements.txt` or other environment details provided.

We have attempted to contact the corresponding authors via email (sent 2021-06-04) for assistance running their code, but received no response at the time of writing.

##### *3.5 MXfold2*

Source code is published, and training is reproducible.

##### *3.6 UFold*

The authors of UFold publish the entire source code on Github, however, we were unable to re-train their network as per their instructions out of the box. After opening an issue on Github[2], the authors were helpful in fixing some issues with the training code. After the authors' help, we were able to reproduce their training methodology, albeit without the use of synthetic sequences [2] for pre-training.

[1] https://github.com/linyuwangPHD/RNA-Secondary-Structure-Database/issues/3
[2] https://github.com/uci-cbcl/UFold/issues/4

#### 4 Mean minimum tree edit distance

##### 4.1 Method

Pair-wise tree-editing distances [3] were calculated using ViennaRNA's [4] RNAdistance[3] program between all sequences in the training and testing sets. For each sequence in the testing set, the minimum tree-editing distance to the sequences in the training set was found. These minimums were then averaged across each cross-validation fold.

##### 4.2 Results

Table 1: Average minimum tree-editing distances of *k*-fold cross-validation training and testing sets. Note the lower distance (i.e. higher structural similarity) compared to family-fold cross-validation sets from Table 2.

| Test set | Train # | Test # | Distance |
|---|---|---|---|
| 0-fold | 2783 | 773 | 16.51 |
| 1-fold | 2783 | 773 | 14.68 |
| 2-fold | 2783 | 773 | 15.17 |
| 3-fold | 2783 | 773 | 16.29 |
| 4-fold | 2783 | 773 | 14.92 |
| Mean | | | 15.51 |

Table 2: Average minimum tree-editing distances of *k*-fold cross-validation training and testing sets. Note the higher distance (i.e. lower structural similarity) compared to *k*-fold cross-validation sets from Table 1.

| Test set | Train # | Test # | Distance |
|---|---|---|---|
| 5s-fold | 2324 | 1283 | 54.50 |
| 16s-fold | 3419 | 67 | 150.03 |
| 23s-fold | 3465 | 15 | 175.60 |
| grp1-fold | 3412 | 74 | 182.08 |
| RNaseP-fold | 3070 | 454 | 161.65 |
| srp-fold | 2653 | 918 | 99.01 |
| telomerase-fold | 3447 | 35 | 185.46 |
| tmRNA-fold | 3063 | 462 | 168.42 |
| tRNA-fold | 2978 | 557 | 38.13 |
| Mean | | | 134.99 |

[3] https://www.tbi.univie.ac.at/RNA/RNAdistance.1.html

#### 5 Demonstrative model results

Table 3: Detailed results of family-fold cross-validation on our demonstrative model.

| | | PPV | | Sensitivity | | $F_1$ | | |
|---|---|---|---|---|---|---|---|---|
| **Family** | **#** | **Baseline** | **Model** | **Baseline** | **Model** | **Baseline** | **Model** | **p-value**[4] |
| 16S rRNA | 67 | 0.54 | 0.47 | 0.62 | 0.50 | 0.58 | 0.48 | <0.001 |
| 23S rRNA | 15 | 0.70 | 0.53 | 0.77 | 0.56 | 0.73 | 0.54 | 0.033 |
| 5S rRNA | 1283 | 0.61 | 0.50 | 0.66 | 0.44 | 0.63 | 0.46 | <0.001 |
| RNase P RNA | 454 | 0.53 | 0.48 | 0.58 | 0.49 | 0.55 | 0.48 | <0.001 |
| Group I Intron | 74 | 0.50 | 0.47 | 0.57 | 0.52 | 0.53 | 0.49 | 0.002 |
| SRP RNA | 918 | 0.62 | 0.51 | 0.67 | 0.51 | 0.64 | 0.50 | <0.001 |
| tRNA | 557 | 0.79 | 0.67 | 0.82 | 0.64 | 0.80 | 0.65 | <0.001 |
| Telomerase RNA | 35 | 0.43 | 0.39 | 0.60 | 0.53 | 0.50 | 0.45 | 0.005 |
| tmRNA | 462 | 0.41 | 0.39 | 0.46 | 0.42 | 0.43 | 0.41 | <0.001 |
| Total | 3865 | | | | | | | |
| Mean | | 0.57 | 0.49 | 0.64 | 0.51 | 0.60 | 0.50 | <0.001 |

Table 4: Detailed results of *k*-fold cross-validation on our demonstrative model.

| | | PPV | | Sensitivity | | $F_1$ | | |
|---|---|---|---|---|---|---|---|---|
| **Family** | **#** | **Baseline** | **Model** | **Baseline** | **Model** | **Baseline** | **Model** | **p-value**[4] |
| 16S rRNA | 67 | 0.54 | 0.58 | 0.62 | 0.61 | 0.58 | 0.60 | 0.482 |
| 23S rRNA | 15 | 0.70 | 0.66 | 0.77 | 0.70 | 0.73 | 0.68 | 0.468 |
| 5S rRNA | 1283 | 0.61 | 0.93 | 0.66 | 0.96 | 0.63 | 0.94 | <0.001 |
| RNase P RNA | 454 | 0.53 | 0.65 | 0.58 | 0.67 | 0.55 | 0.66 | <0.001 |
| Group I Intron | 74 | 0.50 | 0.51 | 0.57 | 0.55 | 0.53 | 0.53 | 0.784 |
| SRP RNA | 918 | 0.62 | 0.80 | 0.67 | 0.83 | 0.64 | 0.81 | <0.001 |
| tRNA | 557 | 0.79 | 0.97 | 0.82 | 0.98 | 0.80 | 0.97 | <0.001 |
| Telomerase RNA | 35 | 0.43 | 0.55 | 0.60 | 0.69 | 0.50 | 0.61 | 0.001 |
| tmRNA | 462 | 0.41 | 0.65 | 0.46 | 0.63 | 0.43 | 0.64 | <0.001 |
| Total | 3865 | | | | | | | |
| Mean | | 0.57 | 0.70 | 0.64 | 0.74 | 0.60 | 0.72 | <0.001 |

Table 5: Area Under the Receiver Operating Characteristic Curve (AUC) for *k*-fold cross-validation and family-fold cross validation.

| | | AUC | | |
|---|---|---|---|---|
| **Family** | **#** | ***k*-fold** | **family-fold** | **p-value** |
| 16S rRNA | 67 | 0.77 | 0.72 | 0.009 |
| 23S rRNA | 15 | 0.79 | 0.73 | 0.080 |
| 5S rRNA | 1283 | 0.95 | 0.72 | <0.001 |
| RNase P RNA | 454 | 0.81 | 0.71 | <0.001 |
| Group I Intron | 74 | 0.73 | 0.72 | 0.387 |
| SRP RNA | 918 | 0.88 | 0.73 | <0.001 |
| tRNA | 557 | 0.97 | 0.79 | <0.001 |
| Telomerase RNA | 35 | 0.76 | 0.68 | <0.001 |
| tmRNA | 462 | 0.82 | 0.68 | <0.001 |
| Total | 3865 | | | |
| Mean | | 0.83 | 0.72 | <0.001 |

[4]Calculated from $F_1$ scores.

#### 6 $\Delta\Delta G'$ nudges

Table 6: $F_1$ score of negative $\Delta\Delta G'$ nudges by family. Red indicates degraded performance, while green indicates improved performance over baseline predictions. Values with * indicate statistically significant results (Table 8).

| $\Delta\Delta G$ | 16S rRNA | 23S rRNA | 5S rRNA | RNase P RNA | Group I Intron RNA | SRP RNA | tRNA | Telomerase RNA | tmRNA |
|---|---|---|---|---|---|---|---|---|---|
| -1.00 | 0.49* | 0.57* | 0.51* | 0.48* | 0.46* | 0.55* | 0.64* | 0.28* | 0.32* |
| -0.98 | 0.49* | 0.58* | 0.51* | 0.48* | 0.46* | 0.55* | 0.64* | 0.28* | 0.33* |
| -0.96 | 0.49* | 0.58* | 0.52* | 0.49* | 0.46* | 0.55* | 0.64* | 0.29* | 0.33* |
| -0.94 | 0.49* | 0.58 | 0.52* | 0.49* | 0.47* | 0.55* | 0.65* | 0.29* | 0.33* |
| -0.92 | 0.49* | 0.58 | 0.52* | 0.49* | 0.47* | 0.55* | 0.65* | 0.29* | 0.33* |
| -0.90 | 0.49* | 0.58 | 0.52* | 0.49* | 0.47* | 0.56* | 0.65* | 0.29* | 0.33* |
| -0.88 | 0.49* | 0.58 | 0.52* | 0.49* | 0.47* | 0.56* | 0.65* | 0.29* | 0.33* |
| -0.86 | 0.49* | 0.59 | 0.52* | 0.49* | 0.47* | 0.56* | 0.66* | 0.30* | 0.33* |
| -0.84 | 0.49* | 0.60 | 0.52* | 0.49* | 0.47* | 0.56* | 0.66* | 0.32* | 0.33* |
| -0.82 | 0.49* | 0.60 | 0.52* | 0.49* | 0.47* | 0.56* | 0.66* | 0.32* | 0.33* |
| -0.80 | 0.50* | 0.60 | 0.52* | 0.50* | 0.47* | 0.56* | 0.66* | 0.32* | 0.33* |
| -0.78 | 0.50* | 0.60 | 0.53* | 0.50* | 0.48* | 0.56* | 0.67* | 0.32* | 0.33* |
| -0.76 | 0.50* | 0.62 | 0.53* | 0.50* | 0.48* | 0.56* | 0.67* | 0.32* | 0.33* |
| -0.74 | 0.50* | 0.62 | 0.53* | 0.50* | 0.48* | 0.57* | 0.67* | 0.33* | 0.34* |
| -0.72 | 0.50* | 0.62 | 0.53* | 0.50* | 0.48* | 0.57* | 0.68* | 0.33* | 0.34* |
| -0.70 | 0.51* | 0.62 | 0.53* | 0.50* | 0.48* | 0.57* | 0.68* | 0.33* | 0.34* |
| -0.68 | 0.51* | 0.62 | 0.53* | 0.50* | 0.48* | 0.57* | 0.68* | 0.33* | 0.34* |
| -0.66 | 0.51* | 0.62 | 0.53* | 0.50* | 0.49* | 0.57* | 0.68* | 0.33* | 0.34* |
| -0.64 | 0.51* | 0.63 | 0.54* | 0.51* | 0.49* | 0.57* | 0.69* | 0.34* | 0.34* |
| -0.62 | 0.52* | 0.63 | 0.54* | 0.51* | 0.49* | 0.58* | 0.69* | 0.34* | 0.35* |
| -0.60 | 0.52* | 0.63 | 0.54* | 0.51* | 0.49* | 0.58* | 0.69* | 0.34* | 0.35* |
| -0.58 | 0.52* | 0.63 | 0.54* | 0.51* | 0.49* | 0.58* | 0.70* | 0.35* | 0.35* |
| -0.56 | 0.52* | 0.63 | 0.54* | 0.51* | 0.49* | 0.58* | 0.70* | 0.35* | 0.35* |
| -0.54 | 0.52* | 0.63 | 0.54* | 0.51* | 0.49* | 0.58* | 0.70* | 0.37* | 0.36* |
| -0.52 | 0.52* | 0.63 | 0.54* | 0.51* | 0.49* | 0.58* | 0.70* | 0.37* | 0.36* |
| -0.50 | 0.52* | 0.64 | 0.54* | 0.51* | 0.49* | 0.59* | 0.71* | 0.37* | 0.36* |
| -0.48 | 0.53* | 0.65 | 0.55* | 0.52* | 0.49* | 0.59* | 0.71* | 0.37* | 0.36* |
| -0.46 | 0.53* | 0.65 | 0.55* | 0.52* | 0.49* | 0.59* | 0.71* | 0.38* | 0.37* |
| -0.44 | 0.53* | 0.67 | 0.55* | 0.52* | 0.49* | 0.59* | 0.72* | 0.38* | 0.37* |
| -0.42 | 0.53* | 0.67 | 0.55* | 0.52* | 0.50* | 0.59* | 0.73* | 0.38* | 0.37* |
| -0.40 | 0.53* | 0.68 | 0.56* | 0.52* | 0.50* | 0.59* | 0.73* | 0.39* | 0.37* |
| -0.38 | 0.53* | 0.68 | 0.56* | 0.52* | 0.50* | 0.59* | 0.74* | 0.39* | 0.37* |
| -0.36 | 0.53* | 0.69 | 0.56* | 0.52* | 0.50* | 0.60* | 0.74* | 0.41* | 0.38* |
| -0.34 | 0.54* | 0.68 | 0.56* | 0.52* | 0.50* | 0.60* | 0.74* | 0.41* | 0.38* |
| -0.32 | 0.54* | 0.71 | 0.57* | 0.52* | 0.50* | 0.60* | 0.74* | 0.42* | 0.38* |
| -0.30 | 0.55 | 0.73 | 0.57* | 0.53* | 0.50* | 0.60* | 0.74* | 0.42* | 0.38* |
| -0.28 | 0.55 | 0.73 | 0.57* | 0.53* | 0.50* | 0.61* | 0.75* | 0.43* | 0.39* |
| -0.26 | 0.55 | 0.73 | 0.57* | 0.53* | 0.51* | 0.61* | 0.75* | 0.44* | 0.39* |
| -0.24 | 0.56 | 0.73 | 0.58* | 0.53* | 0.51* | 0.61* | 0.75* | 0.45* | 0.39* |
| -0.22 | 0.56 | 0.73 | 0.58* | 0.53* | 0.51* | 0.62* | 0.76* | 0.45* | 0.40* |
| -0.20 | 0.56 | 0.73 | 0.58* | 0.53* | 0.51* | 0.62* | 0.76* | 0.46 | 0.40* |
| -0.18 | 0.56 | 0.73 | 0.59* | 0.54* | 0.51* | 0.62* | 0.77* | 0.47 | 0.40* |
| -0.16 | 0.56 | 0.72 | 0.59* | 0.54* | 0.52 | 0.62* | 0.77* | 0.48 | 0.40* |
| -0.14 | 0.56 | 0.72 | 0.60* | 0.54* | 0.52 | 0.63 | 0.78* | 0.47 | 0.41* |
| -0.12 | 0.56 | 0.72 | 0.60* | 0.54* | 0.52 | 0.64 | 0.78* | 0.47* | 0.41* |
| -0.10 | 0.56 | 0.74 | 0.61* | 0.54* | 0.53 | 0.64 | 0.78* | 0.47* | 0.41* |
| -0.08 | 0.56 | 0.74 | 0.61* | 0.54* | 0.52 | 0.64 | 0.78* | 0.48 | 0.42* |
| -0.06 | 0.57 | 0.74 | 0.62* | 0.55 | 0.52* | 0.64 | 0.79* | 0.49 | 0.42* |
| -0.04 | 0.56 | 0.72 | 0.62* | 0.55 | 0.53 | 0.64 | 0.80* | 0.49 | 0.42* |
| -0.02 | 0.56 | 0.72 | 0.63* | 0.55 | 0.53 | 0.63 | 0.80 | 0.49 | 0.43* |
| 0.0 | 0.57 | 0.73 | 0.63 | 0.55 | 0.53 | 0.63 | 0.80 | 0.50 | 0.43 |

Table 7: $F_1$ score of positive $\Delta\Delta G'$ nudges by family. Red indicates degraded performance, while green indicates improved performance over baseline predictions. Values with * indicate statistically significant results (Table 9).

| $\Delta\Delta G$ | 16S rRNA | 23S rRNA | 5S rRNA | RNase P RNA | Group I Intron RNA | SRP RNA | tRNA | Telomerase RNA | tmRNA |
|---|---|---|---|---|---|---|---|---|---|
| 0.02 | 0.57 | 0.73 | 0.63 | 0.55 | 0.54 | 0.63 | 0.81 | 0.50 | 0.44* |
| 0.04 | 0.57 | 0.72 | 0.63 | 0.55 | 0.53 | 0.63 | 0.81* | 0.51 | 0.44* |
| 0.06 | 0.56* | 0.66 | 0.65* | 0.55 | 0.53 | 0.63 | 0.80 | 0.52 | 0.45* |
| 0.08 | 0.56 | 0.66 | 0.65* | 0.56 | 0.53 | 0.62* | 0.80 | 0.53 | 0.45* |
| 0.10 | 0.56 | 0.68 | 0.65* | 0.55 | 0.53 | 0.61* | 0.80 | 0.53 | 0.46* |
| 0.12 | 0.56 | 0.67 | 0.65* | 0.55 | 0.54 | 0.60* | 0.81 | 0.53 | 0.46* |
| 0.14 | 0.55 | 0.67 | 0.64* | 0.55 | 0.54 | 0.60* | 0.81 | 0.53 | 0.46* |
| 0.16 | 0.54* | 0.68 | 0.64 | 0.54* | 0.53 | 0.59* | 0.80 | 0.52 | 0.46* |
| 0.18 | 0.53* | 0.66 | 0.62 | 0.54* | 0.53 | 0.58* | 0.79 | 0.52 | 0.46* |
| 0.20 | 0.54* | 0.66 | 0.61* | 0.53* | 0.53 | 0.56* | 0.78* | 0.51 | 0.46* |
| 0.22 | 0.52* | 0.67 | 0.60* | 0.53* | 0.49* | 0.55* | 0.76* | 0.51 | 0.45* |
| 0.24 | 0.50* | 0.66 | 0.58* | 0.52* | 0.47* | 0.54* | 0.76* | 0.49 | 0.44 |
| 0.26 | 0.49* | 0.65 | 0.56* | 0.50* | 0.46* | 0.51* | 0.74* | 0.47 | 0.44 |
| 0.28 | 0.48* | 0.63 | 0.53* | 0.49* | 0.44* | 0.47* | 0.73* | 0.45 | 0.43 |
| 0.30 | 0.46* | 0.57* | 0.51* | 0.47* | 0.41* | 0.44* | 0.70* | 0.42* | 0.43 |
| 0.32 | 0.45* | 0.54* | 0.47* | 0.46* | 0.38* | 0.40* | 0.67* | 0.42* | 0.41* |
| 0.34 | 0.44* | 0.50* | 0.44* | 0.44* | 0.36* | 0.36* | 0.62* | 0.41* | 0.39* |
| 0.36 | 0.40* | 0.47* | 0.41* | 0.43* | 0.32* | 0.34* | 0.58* | 0.38* | 0.38* |
| 0.38 | 0.37* | 0.45* | 0.36* | 0.41* | 0.28* | 0.31* | 0.54* | 0.36* | 0.34* |
| 0.40 | 0.34* | 0.41* | 0.32* | 0.38* | 0.25* | 0.27* | 0.49* | 0.33* | 0.31* |
| 0.42 | 0.32* | 0.33* | 0.28* | 0.36* | 0.23* | 0.25* | 0.44* | 0.30* | 0.29* |
| 0.44 | 0.29* | 0.30* | 0.24* | 0.33* | 0.21* | 0.19* | 0.41* | 0.26* | 0.25* |
| 0.46 | 0.29* | 0.25* | 0.21* | 0.30* | 0.19* | 0.16* | 0.34* | 0.23* | 0.22* |
| 0.48 | 0.27* | 0.22* | 0.16* | 0.27* | 0.15* | 0.13* | 0.26* | 0.21* | 0.20* |
| 0.50 | 0.22* | 0.21* | 0.13* | 0.24* | 0.12* | 0.10* | 0.22* | 0.20* | 0.16* |
| 0.52 | 0.19* | 0.17* | 0.10* | 0.22* | 0.10* | 0.09* | 0.17* | 0.17* | 0.13* |
| 0.54 | 0.16* | 0.15* | 0.07* | 0.19* | 0.08* | 0.08* | 0.12* | 0.16* | 0.11* |
| 0.56 | 0.15* | 0.10* | 0.05* | 0.17* | 0.07* | 0.07* | 0.10* | 0.13* | 0.10* |
| 0.58 | 0.15* | 0.09* | 0.04* | 0.15* | 0.06* | 0.06* | 0.08* | 0.12* | 0.08* |
| 0.60 | 0.11* | 0.07* | 0.03* | 0.11* | 0.04* | 0.05* | 0.06* | 0.06* | 0.06* |
| 0.62 | 0.09* | 0.06* | 0.02* | 0.09* | 0.03* | 0.04* | 0.04* | 0.05* | 0.05* |
| 0.64 | 0.06* | 0.05* | 0.02* | 0.07* | 0.03* | 0.02* | 0.04* | 0.03* | 0.04* |
| 0.66 | 0.06* | 0.05* | 0.01* | 0.06* | 0.01* | 0.02* | 0.03* | 0.03* | 0.04* |
| 0.68 | 0.05* | 0.03* | 0.01* | 0.05* | 0.01* | 0.02* | 0.03* | 0.02* | 0.03* |
| 0.70 | 0.05* | 0.03* | 0.01* | 0.04* | 0.01* | 0.01* | 0.02* | 0.02* | 0.02* |
| 0.72 | 0.05* | 0.03* | 0.01* | 0.03* | 0.01* | 0.01* | 0.02* | 0.01* | 0.02* |
| 0.74 | 0.05* | 0.01* | 0.01* | 0.03* | 0.01* | 0.01* | 0.02* | 0.01* | 0.02* |
| 0.76 | 0.05* | 0.01* | 0.01* | 0.02* | 0.01* | 0.01* | 0.02* | 0.01* | 0.02* |
| 0.78 | 0.05* | 0.01* | 0.01* | 0.02* | 0.01* | 0.01* | 0.02* | 0.01* | 0.02* |
| 0.80 | 0.05* | 0.01* | 0.01* | 0.02* | 0.01* | 0.01* | 0.02* | 0.01* | 0.02* |
| 0.82 | 0.05* | 0.01* | 0.01* | 0.02* | 0.01* | 0.01* | 0.02* | 0.01* | 0.02* |
| 0.84 | 0.05* | 0.01* | 0.01* | 0.02* | 0.01* | 0.01* | 0.02* | 0.02* | 0.02* |
| 0.86 | 0.05* | 0.01* | 0.01* | 0.02* | 0.01* | 0.01* | 0.02* | 0.02* | 0.02* |
| 0.88 | 0.04* | 0.00* | 0.01* | 0.02* | 0.01* | 0.01* | 0.02* | 0.02* | 0.02* |
| 0.90 | 0.04* | 0.00* | 0.01* | 0.02* | 0.01* | 0.01* | 0.02* | 0.02* | 0.02* |
| 0.92 | 0.04* | 0.00* | 0.01* | 0.02* | 0.01* | 0.01* | 0.02* | 0.02* | 0.02* |
| 0.94 | 0.03* | 0.00* | 0.01* | 0.02* | 0.01* | 0.01* | 0.02* | 0.01* | 0.02* |
| 0.96 | 0.03* | 0.00* | 0.01* | 0.02* | 0.01* | 0.01* | 0.02* | 0.02* | 0.02* |
| 0.98 | 0.03* | 0.00* | 0.01* | 0.02* | 0.01* | 0.01* | 0.02* | 0.02* | 0.02* |
| 1.00 | 0.03* | 0.00* | 0.01* | 0.02* | 0.01* | 0.01* | 0.02* | 0.02* | 0.02* |
| 0.0 | 0.57 | 0.73 | 0.63 | 0.55 | 0.53 | 0.63 | 0.80 | 0.50 | 0.43 |

Table 8: Two-tailed paired t-tests for positive $\Delta\Delta G'$ nudges.

| $\Delta\Delta G$ | 16S rRNA | 23S rRNA | 5S rRNA | RNase P RNA | Group I Intron RNA | SRP RNA | tRNA | Telomerase RNA | tmRNA |
|---|---|---|---|---|---|---|---|---|---|
| -1.00 | <0.001 | 0.036 | <0.001 | <0.001 | <0.001 | <0.001 | <0.001 | <0.001 | <0.001 |
| -0.98 | <0.001 | 0.037 | <0.001 | <0.001 | <0.001 | <0.001 | <0.001 | <0.001 | <0.001 |
| -0.96 | <0.001 | 0.037 | <0.001 | <0.001 | <0.001 | <0.001 | <0.001 | <0.001 | <0.001 |
| -0.94 | <0.001 | 0.051 | <0.001 | <0.001 | <0.001 | <0.001 | <0.001 | <0.001 | <0.001 |
| -0.92 | <0.001 | 0.051 | <0.001 | <0.001 | <0.001 | <0.001 | <0.001 | <0.001 | <0.001 |
| -0.90 | <0.001 | 0.052 | <0.001 | <0.001 | <0.001 | <0.001 | <0.001 | <0.001 | <0.001 |
| -0.88 | <0.001 | 0.052 | <0.001 | <0.001 | 0.001 | <0.001 | <0.001 | <0.001 | <0.001 |
| -0.86 | <0.001 | 0.062 | <0.001 | <0.001 | 0.001 | <0.001 | <0.001 | <0.001 | <0.001 |
| -0.84 | <0.001 | 0.055 | <0.001 | <0.001 | <0.001 | <0.001 | <0.001 | <0.001 | <0.001 |
| -0.82 | <0.001 | 0.055 | <0.001 | <0.001 | <0.001 | <0.001 | <0.001 | <0.001 | <0.001 |
| -0.80 | <0.001 | 0.055 | <0.001 | <0.001 | <0.001 | <0.001 | <0.001 | <0.001 | <0.001 |
| -0.78 | <0.001 | 0.055 | <0.001 | <0.001 | 0.001 | <0.001 | <0.001 | <0.001 | <0.001 |
| -0.76 | <0.001 | 0.076 | <0.001 | <0.001 | <0.001 | <0.001 | <0.001 | <0.001 | <0.001 |
| -0.74 | <0.001 | 0.076 | <0.001 | <0.001 | 0.001 | <0.001 | <0.001 | <0.001 | <0.001 |
| -0.72 | <0.001 | 0.083 | <0.001 | <0.001 | 0.001 | <0.001 | <0.001 | <0.001 | <0.001 |
| -0.70 | 0.001 | 0.083 | <0.001 | <0.001 | <0.001 | <0.001 | <0.001 | <0.001 | <0.001 |
| -0.68 | 0.001 | 0.083 | <0.001 | <0.001 | <0.001 | <0.001 | <0.001 | <0.001 | <0.001 |
| -0.66 | 0.001 | 0.085 | <0.001 | <0.001 | 0.001 | <0.001 | <0.001 | <0.001 | <0.001 |
| -0.64 | 0.002 | 0.097 | <0.001 | <0.001 | 0.001 | <0.001 | <0.001 | <0.001 | <0.001 |
| -0.62 | 0.002 | 0.103 | <0.001 | <0.001 | 0.001 | <0.001 | <0.001 | <0.001 | <0.001 |
| -0.60 | 0.002 | 0.103 | <0.001 | <0.001 | 0.001 | <0.001 | <0.001 | <0.001 | <0.001 |
| -0.58 | 0.004 | 0.099 | <0.001 | <0.001 | 0.001 | <0.001 | <0.001 | <0.001 | <0.001 |
| -0.56 | 0.005 | 0.099 | <0.001 | <0.001 | 0.001 | <0.001 | <0.001 | <0.001 | <0.001 |
| -0.54 | 0.005 | 0.089 | <0.001 | <0.001 | 0.001 | <0.001 | <0.001 | <0.001 | <0.001 |
| -0.52 | 0.005 | 0.090 | <0.001 | <0.001 | 0.001 | <0.001 | <0.001 | <0.001 | <0.001 |
| -0.50 | 0.006 | 0.129 | <0.001 | <0.001 | 0.001 | <0.001 | <0.001 | <0.001 | <0.001 |
| -0.48 | 0.011 | 0.134 | <0.001 | <0.001 | 0.001 | <0.001 | <0.001 | <0.001 | <0.001 |
| -0.46 | 0.014 | 0.132 | <0.001 | <0.001 | 0.001 | <0.001 | <0.001 | <0.001 | <0.001 |
| -0.44 | 0.013 | 0.214 | <0.001 | <0.001 | 0.002 | <0.001 | <0.001 | <0.001 | <0.001 |
| -0.42 | 0.010 | 0.214 | <0.001 | <0.001 | 0.003 | <0.001 | <0.001 | <0.001 | <0.001 |
| -0.40 | 0.012 | 0.330 | <0.001 | <0.001 | 0.004 | <0.001 | <0.001 | 0.001 | <0.001 |
| -0.38 | 0.018 | 0.336 | <0.001 | <0.001 | 0.006 | <0.001 | <0.001 | 0.001 | <0.001 |
| -0.36 | 0.018 | 0.377 | <0.001 | <0.001 | 0.006 | <0.001 | <0.001 | 0.002 | <0.001 |
| -0.34 | 0.032 | 0.369 | <0.001 | <0.001 | 0.006 | <0.001 | <0.001 | 0.002 | <0.001 |
| -0.32 | 0.032 | 0.615 | <0.001 | <0.001 | 0.006 | <0.001 | <0.001 | 0.004 | <0.001 |
| -0.03 | 0.057 | 0.935 | <0.001 | <0.001 | 0.005 | <0.001 | <0.001 | 0.005 | <0.001 |
| -0.28 | 0.057 | 0.935 | <0.001 | <0.001 | 0.008 | <0.001 | <0.001 | 0.010 | <0.001 |
| -0.26 | 0.058 | 0.935 | <0.001 | <0.001 | 0.012 | <0.001 | <0.001 | 0.016 | <0.001 |
| -0.24 | 0.155 | 0.935 | <0.001 | <0.001 | 0.034 | <0.001 | <0.001 | 0.042 | <0.001 |
| -0.22 | 0.189 | 0.938 | <0.001 | <0.001 | 0.014 | <0.001 | <0.001 | 0.043 | <0.001 |
| -0.20 | 0.209 | 0.943 | <0.001 | <0.001 | 0.019 | <0.001 | <0.001 | 0.065 | <0.001 |
| -0.18 | 0.201 | 0.909 | <0.001 | <0.001 | 0.042 | <0.001 | <0.001 | 0.141 | <0.001 |
| -0.16 | 0.244 | 0.687 | <0.001 | 0.003 | 0.080 | 0.002 | <0.001 | 0.224 | <0.001 |
| -0.14 | 0.186 | 0.809 | <0.001 | 0.008 | 0.099 | 0.275 | <0.001 | 0.133 | <0.001 |
| -0.12 | 0.284 | 0.775 | <0.001 | 0.008 | 0.080 | 0.394 | <0.001 | 0.019 | <0.001 |
| -0.10 | 0.217 | 0.815 | <0.001 | 0.001 | 0.211 | 0.430 | <0.001 | 0.014 | <0.001 |
| -0.08 | 0.245 | 0.801 | <0.001 | 0.007 | 0.117 | 0.347 | <0.001 | 0.178 | <0.001 |
| -0.06 | 0.624 | 0.709 | <0.001 | 0.318 | 0.039 | 0.095 | 0.017 | 0.279 | <0.001 |
| -0.04 | 0.303 | 0.243 | 0.004 | 0.321 | 0.317 | 0.092 | 0.046 | 0.200 | <0.001 |
| -0.02 | 0.209 | 0.308 | 0.006 | 0.749 | 0.491 | 0.654 | 0.054 | 0.233 | <0.001 |

Table 9: Two-tailed paired t-tests for negative $\Delta\Delta G'$ nudges.

| $\Delta\Delta G$ | 16S rRNA | 23S rRNA | 5S rRNA | RNase P RNA | Group I Intron RNA | SRP RNA | tRNA | Telomerase RNA | tmRNA |
|---|---|---|---|---|---|---|---|---|---|
| 0.02 | 0.548 | 0.102 | 0.193 | 0.363 | 0.122 | 0.669 | 0.143 | 0.900 | 0.009 |
| 0.04 | 0.146 | 0.432 | 0.571 | 0.828 | 0.406 | 0.684 | 0.034 | 0.423 | 0.001 |
| 0.06 | 0.042 | 0.213 | <0.001 | 0.652 | 0.276 | 0.181 | 0.643 | 0.318 | <0.001 |
| 0.08 | 0.088 | 0.213 | <0.001 | 0.408 | 0.383 | 0.007 | 0.733 | 0.121 | <0.001 |
| 0.10 | 0.134 | 0.372 | <0.001 | 0.825 | 0.631 | <0.001 | 0.626 | 0.113 | <0.001 |
| 0.12 | 0.117 | 0.327 | 0.001 | 0.651 | 0.864 | <0.001 | 0.323 | 0.105 | <0.001 |
| 0.14 | 0.074 | 0.345 | 0.013 | 0.181 | 0.697 | <0.001 | 0.467 | 0.164 | <0.001 |
| 0.16 | 0.044 | 0.379 | 0.350 | 0.046 | 0.812 | <0.001 | 0.983 | 0.312 | <0.001 |
| 0.18 | 0.011 | 0.269 | 0.121 | 0.016 | 0.512 | <0.001 | 0.090 | 0.301 | <0.001 |
| 0.20 | 0.031 | 0.257 | <0.001 | 0.001 | 0.783 | <0.001 | 0.025 | 0.603 | <0.001 |
| 0.22 | 0.006 | 0.401 | <0.001 | <0.001 | 0.013 | <0.001 | <0.001 | 0.762 | 0.002 |
| 0.24 | 0.001 | 0.292 | <0.001 | <0.001 | 0.001 | <0.001 | <0.001 | 0.824 | 0.070 |
| 0.26 | <0.001 | 0.198 | <0.001 | <0.001 | <0.001 | <0.001 | <0.001 | 0.229 | 0.432 |
| 0.28 | <0.001 | 0.124 | <0.001 | <0.001 | <0.001 | <0.001 | <0.001 | 0.083 | 0.888 |
| 0.30 | <0.001 | 0.011 | <0.001 | <0.001 | <0.001 | <0.001 | <0.001 | 0.012 | 0.354 |
| 0.32 | <0.001 | 0.004 | <0.001 | <0.001 | <0.001 | <0.001 | <0.001 | 0.012 | 0.019 |
| 0.34 | <0.001 | 0.002 | <0.001 | <0.001 | <0.001 | <0.001 | <0.001 | 0.004 | <0.001 |
| 0.36 | <0.001 | 0.001 | <0.001 | <0.001 | <0.001 | <0.001 | <0.001 | 0.001 | <0.001 |
| 0.38 | <0.001 | 0.001 | <0.001 | <0.001 | <0.001 | <0.001 | <0.001 | <0.001 | <0.001 |
| 0.40 | <0.001 | <0.001 | <0.001 | <0.001 | <0.001 | <0.001 | <0.001 | <0.001 | <0.001 |
| 0.42 | <0.001 | <0.001 | <0.001 | <0.001 | <0.001 | <0.001 | <0.001 | <0.001 | <0.001 |
| 0.44 | <0.001 | <0.001 | <0.001 | <0.001 | <0.001 | <0.001 | <0.001 | <0.001 | <0.001 |
| 0.46 | <0.001 | <0.001 | <0.001 | <0.001 | <0.001 | <0.001 | <0.001 | <0.001 | <0.001 |
| 0.48 | <0.001 | <0.001 | <0.001 | <0.001 | <0.001 | <0.001 | <0.001 | <0.001 | <0.001 |
| 0.50 | <0.001 | <0.001 | <0.001 | <0.001 | <0.001 | <0.001 | <0.001 | <0.001 | <0.001 |
| 0.52 | <0.001 | <0.001 | <0.001 | <0.001 | <0.001 | <0.001 | <0.001 | <0.001 | <0.001 |
| 0.54 | <0.001 | <0.001 | <0.001 | <0.001 | <0.001 | <0.001 | <0.001 | <0.001 | <0.001 |
| 0.56 | <0.001 | <0.001 | <0.001 | <0.001 | <0.001 | <0.001 | <0.001 | <0.001 | <0.001 |
| 0.58 | <0.001 | <0.001 | <0.001 | <0.001 | <0.001 | <0.001 | <0.001 | <0.001 | <0.001 |
| 0.60 | <0.001 | <0.001 | <0.001 | <0.001 | <0.001 | <0.001 | <0.001 | <0.001 | <0.001 |
| 0.62 | <0.001 | <0.001 | <0.001 | <0.001 | <0.001 | <0.001 | <0.001 | <0.001 | <0.001 |
| 0.64 | <0.001 | <0.001 | <0.001 | <0.001 | <0.001 | <0.001 | <0.001 | <0.001 | <0.001 |
| 0.66 | <0.001 | <0.001 | <0.001 | <0.001 | <0.001 | <0.001 | <0.001 | <0.001 | <0.001 |
| 0.68 | <0.001 | <0.001 | <0.001 | <0.001 | <0.001 | <0.001 | <0.001 | <0.001 | <0.001 |
| 0.70 | <0.001 | <0.001 | <0.001 | <0.001 | <0.001 | <0.001 | <0.001 | <0.001 | <0.001 |
| 0.72 | <0.001 | <0.001 | <0.001 | <0.001 | <0.001 | <0.001 | <0.001 | <0.001 | <0.001 |
| 0.74 | <0.001 | <0.001 | <0.001 | <0.001 | <0.001 | <0.001 | <0.001 | <0.001 | <0.001 |
| 0.76 | <0.001 | <0.001 | <0.001 | <0.001 | <0.001 | <0.001 | <0.001 | <0.001 | <0.001 |
| 0.78 | <0.001 | <0.001 | <0.001 | <0.001 | <0.001 | <0.001 | <0.001 | <0.001 | <0.001 |
| 0.80 | <0.001 | <0.001 | <0.001 | <0.001 | <0.001 | <0.001 | <0.001 | <0.001 | <0.001 |
| 0.82 | <0.001 | <0.001 | <0.001 | <0.001 | <0.001 | <0.001 | <0.001 | <0.001 | <0.001 |
| 0.84 | <0.001 | <0.001 | <0.001 | <0.001 | <0.001 | <0.001 | <0.001 | <0.001 | <0.001 |
| 0.86 | <0.001 | <0.001 | <0.001 | <0.001 | <0.001 | <0.001 | <0.001 | <0.001 | <0.001 |
| 0.88 | <0.001 | <0.001 | <0.001 | <0.001 | <0.001 | <0.001 | <0.001 | <0.001 | <0.001 |
| 0.90 | <0.001 | <0.001 | <0.001 | <0.001 | <0.001 | <0.001 | <0.001 | <0.001 | <0.001 |
| 0.92 | <0.001 | <0.001 | <0.001 | <0.001 | <0.001 | <0.001 | <0.001 | <0.001 | <0.001 |
| 0.94 | <0.001 | <0.001 | <0.001 | <0.001 | <0.001 | <0.001 | <0.001 | <0.001 | <0.001 |
| 0.96 | <0.001 | <0.001 | <0.001 | <0.001 | <0.001 | <0.001 | <0.001 | <0.001 | <0.001 |
| 0.98 | <0.001 | <0.001 | <0.001 | <0.001 | <0.001 | <0.001 | <0.001 | <0.001 | <0.001 |
| 1.00 | <0.001 | <0.001 | <0.001 | <0.001 | <0.001 | <0.001 | <0.001 | <0.001 | <0.001 |
